# Anticipatory smooth pursuit eye movements scale with the probability of visual motion: the role of target speed and acceleration

**DOI:** 10.1101/2023.10.31.564614

**Authors:** Vanessa Carneiro Morita, David Souto, Guillaume S. Masson, Anna Montagnini

## Abstract

Sensory-motor systems can extract statistical regularities in dynamic uncertain environments, enabling quicker responses and anticipatory behavior for expected events. Anticipatory smooth pursuit eye movements (aSP) have been observed in primates when the temporal and kinematic properties of a forthcoming visual moving target are fully or partially predictable. To investigate the nature of the internal model of target kinematics underlying aSP, we tested the effect of varying the target kinematics and its predictability. Participants tracked a small visual target in a constant direction with either constant, accelerating or decelerating speed. Across experimental blocks, we manipulated the probability of each kinematic condition varying either speed or acceleration across trials; with either one kinematic condition (providing certainty) or with a mixture of conditions with a fixed probability within a block. We show that aSP is robustly modulated by target kinematics. With constant-velocity targets, aSP velocity scales linearly with target velocity in blocked sessions, and matches the probability-weighted average in the mixture sessions. Predictable target acceleration does also have an influence on aSP, suggesting that the internal model of motion which drives anticipation contains some information about the changing target kinematics, beyond the initial target speed. However, there is a large variability across participants in the precision and consistency with which this information is taken into account in order to control anticipatory behavior.

## Introduction

Smooth pursuit eye movements allow us to maintain the image of a moving object of interest steady on the retinas. To do so, tracking eye movements rely primarily on the neural representation of the moving target’s speed and direction (Carl & Gellman, 1987; Lisberger & Westbrook,; Tychsen & Lisberger, 1986). When the target moves at constant speed across the visual field, the eyes start accelerating in the same direction as the target motion with a short latency (∼100-130 ms in humans; Carl & Gellman, 1987). In the optimal speed range for human pursuit (i.e. below 20-30°/s), the eyes typically reach a steady state velocity close to the target’s velocity within ∼300 ms from visual motion onset. Steady-state smooth tracking, in close coordination with the so-called catch-up saccades can dynamically maintain a good alignment between the fovea and the target retinal image position (Carl & Gellman, 1987; Orban De Xivry & Lefèvre, 2007). Even though natural objects rarely move at a constant velocity, only a few studies have investigated eye tracking behavior for accelerating (or decelerating) targets. Those studies reported that humans are indeed able to track visible targets with smoothly-varying speed (e.g. Bennett & Benguigui, 2013; Kreyenmeier et al., 2022), but with weak accuracy. Like perceptual discrimination judgments, tracking eye movements discriminate poorly between target accelerations as compared to target speeds in humans (e.g., Watamaniuk & Heinen, 2003; Kowler & McKee, 1987) and macaque monkeys (Lisberger & Movshon, 1999).

Most of the early work on pursuit eye movements and visual speed/acceleration processing were concerned with the online control. However, our dynamic environment is often, at least in part, predictable, because either the motion is produced by our own body (e.g., Gauthier et al., 1988; Landelle et al., 2016), or it can be inferred from general prior knowledge, past experience or perceptual cues about an object’s motion (Kowler et al., 2019). For instance, when a target’s velocity changes in a periodic way (e.g., sinusoidal motion), ocular tracking can rapidly take advantage of this predictable motion such that, after only a few cycles, the fovea is aligned to the target position with nearly no lag (Kowler & Steinman, 1979a). Moreover, motion predictability allows anticipation of the target motion onset, or target reappearance after a transient occlusion (e.g. Dodge et al., 1930; for a review, see Kowler et al., 2019 and Fukushima et al., 2013). These anticipatory smooth pursuit eye movements (aSP) are thought to help in quickly reducing the retinal position and velocity errors during the early phase of visually-guided pursuit (e.g. Kao & Morrow, 1994).

Large efforts have been devoted to understanding what kind of signals drive anticipatory responses (Kowler et al., 2019). The underlying computational mechanisms of aSP are however still not fully understood. Empirically, target motion predictability can be manipulated across different time scales (e.g. across or within trials in a standard visuomotor experiment) and repetition schedules. Our group, and others have previously shown that aSP amplitude is proportional to the probability of a given target motion direction in a direction-biased task (Damasse et al., 2018; Montagnini et al., 2010; Rubinstein et al., 2024; Santos & Kowler, 2017; Wu et al., 2021). Expectation for a given target speed also modulates aSP: anticipatory eye velocity increases with speed predictability, in a random versus fully predictable speed design (e.g., Heinen et al., 2005; Jarrett & Barnes, 2002; Kao & Morrow, 1994). aSP also depends on the recent trial history of target speed (Maus et al. 2015). However, it has not been tested whether aSP exhibits the same linear dependency upon speed probability as it does for direction probability.

Predictive aSP has been extensively investigated for simple target trajectories of constant speed and/or direction (e.g., among many others, in Barnes & Asselman, 1991; Heinen et al., 2005; Kowler & Steinman, 1979b). However, much less is known about aSP for accelerating targets and the contribution of acceleration signals to the internal representation of motion raises several questions. First, it is still not known if acceleration is taken into account when preparing for anticipatory movements, or if the latter are based on simple estimates approximating the target motion profile (e.g. instantaneous at some critical moments, or time-averaged speed, Bennett, Orban De Xivry, et al., 2010). Second, most of the previous studies have investigated acceleration-based predictive behavior only on a short time scale, namely during the transient occlusion of a moving target with an accelerating profile (Bennett et al., 2007, 2010; Bennett & Barnes, 2006).

From these previous studies, target acceleration does not appear to be fully integrated in predictive pursuit. For instance, when an accelerating target is occluded after a short presentation of accelerating motion (∼ 200 ms), eye velocity during the blanking period reduces, and its predictive reacceleration prior to the target reappearance is not influenced by the target acceleration (Bennett et al., 2007). However, if the target visibility window before the occlusion is long enough (> 500 ms), eye velocity reduces during the blanking period, but recovers in an acceleration-scaled manner prior to the target reappearance (Bennett, Orban De Xivry, et al., 2010). Moreover, target acceleration is not appropriately taken into account neither in a task where participants are asked to track a target and, after a period of occlusion, to predict the target position, or to predict when the target is going to reach a certain position (Bennett & Benguigui, 2013; Kreyenmeier et al., 2022). However, one of the very few studies addressing the effect of acceleration predictability on a longer time scale (blocked design) showed that the amplitude of aSP observed before the target reappearance in an occlusion paradigm scales with target acceleration (Bennett & Barnes, 2006). Here, we aimed at better understanding how target motion acceleration shapes aSP when the predictability of the kinematic profile is manipulated on a relatively long timescale, namely across blocks of several tenths of trials.

In the present study we tested the following hypotheses: 1) Anticipatory smooth pursuit eye velocity scales linearly with target speed probability (similar to direction probability, Damasse et al. 2018, Santos et al. 2017), across experimental blocks with fixed probability; 2) Anticipatory eye velocity is modulated by target acceleration and 3) it scales with the probability of the accelerating motion. To test these hypotheses, we analyzed anticipatory oculomotor behavior for targets moving in a fully predictable direction but with different speeds and accelerations. We manipulated the probability of each kinematic condition across blocks. We showed that target speed probability influences aSP similarly to motion direction probability, with a linear dependence of aSP velocity upon target speed probability. We also showed that aSP can be driven by predictable accelerating targets, in a way that accounts for the expected change in target velocity and for its probability, even though with large inter-individual variability.

## Methods

### Participants

Twenty-nine healthy adult volunteers signed an informed consent to participate in the experiments presented in this study. The experimental protocol was approved by the Ethics Committee *Comité de Protection des Personnes OUEST III* (CPP reference: PredictEye-2018-A02608-47), in full respect of the Helsinki declaration guidelines. Three of the authors (AM, GM, DS) participated in Experiment 1A (n=3), two of the authors (AM, VCM) participated in Experiment 1B-2A (n=13) and one of the authors (VCM) participated in Experiment 2B (n=5) and in Experiment 3 (n=8). In Experiment 1A, anticipatory eye movements and initial pursuit were recorded with high precision by using the scleral search coil technique (Robinson, 1963) in a small participant sample. A preliminary version of the results from Experiment 1A was presented previously at the VSS conference (Souto et al., 2008). The core finding of this experiment motivated Experiments 1B-2A-2B, that were run with a larger sample of participants, a smaller subset of probability conditions, and using a less invasive technique (video eye tracking). Experiment 3 was designed to test the effect of acceleration on anticipatory eye speed in fully predictable blocks, again using a non-invasive eye recording technique.

### Stimuli and procedure

In all experiments, participants were instructed to visually track a small moving target by smoothly pursuing it with their eyes, as accurately as they could, while their eye movements were recorded. We used different materials across experiments.

#### Experiment 1A

The detailed methods are described elsewhere (Wallace et al. 2005). Briefly, a PC running the REX package controlled both stimulus presentation and data acquisition. Stimuli were generated with an SGI Fuel workstation (ABC Corp., New York, USA, no longer available) and back-projected onto a large translucent screen (80° x 60°, viewing distance: 1m) using a Barco 908s video-projector (1280’’, 1024 pixels at 76 Hz). Oculomotor recordings were collected using the scleral search coil technique (Collewijn et al. 1975).

This experiment probed aSP in different target speed probability contexts (**Figure 1a**). Each trial started with a white fixation point, located at the center of the screen for a random duration between 300 and 450 ms, on a black uniform background (luminance <1 cd/m^2^). If the participant fixated accurately (i.e. within a 2°-side, square electronic window) during the last 200 ms, the fixation target was extinguished and followed by a fixed-duration, 300-ms empty screen. At the end of this gap, the target (a white, gaussian-windowed circle, 0.2° std, maximum luminance 45 cd/m^2^) appeared at the center of the screen and started moving horizontally to the right, for a fixed period of 500 ms. The target speed was either 5.5 °/s (low speed, LS) or 16.5 °/s (high speed, HS). In each experimental block, a different target speed-probability condition defined the proportion of trials at high speed (P(HS)=0, 0.1, 0.25, 0.5, 0.75, 0.9 and 1). The complementary proportion of trials had a low-speed target motion (P(LS)=1-P(HS)). Participants completed 500 trials per block, except for the P(HS)=0 and P(HS)=1 conditions where only 250 trials were completed. One or two blocks were completed in a day, with the constraint of not exceeding a total of one hour of duration.

**Figure 1.**
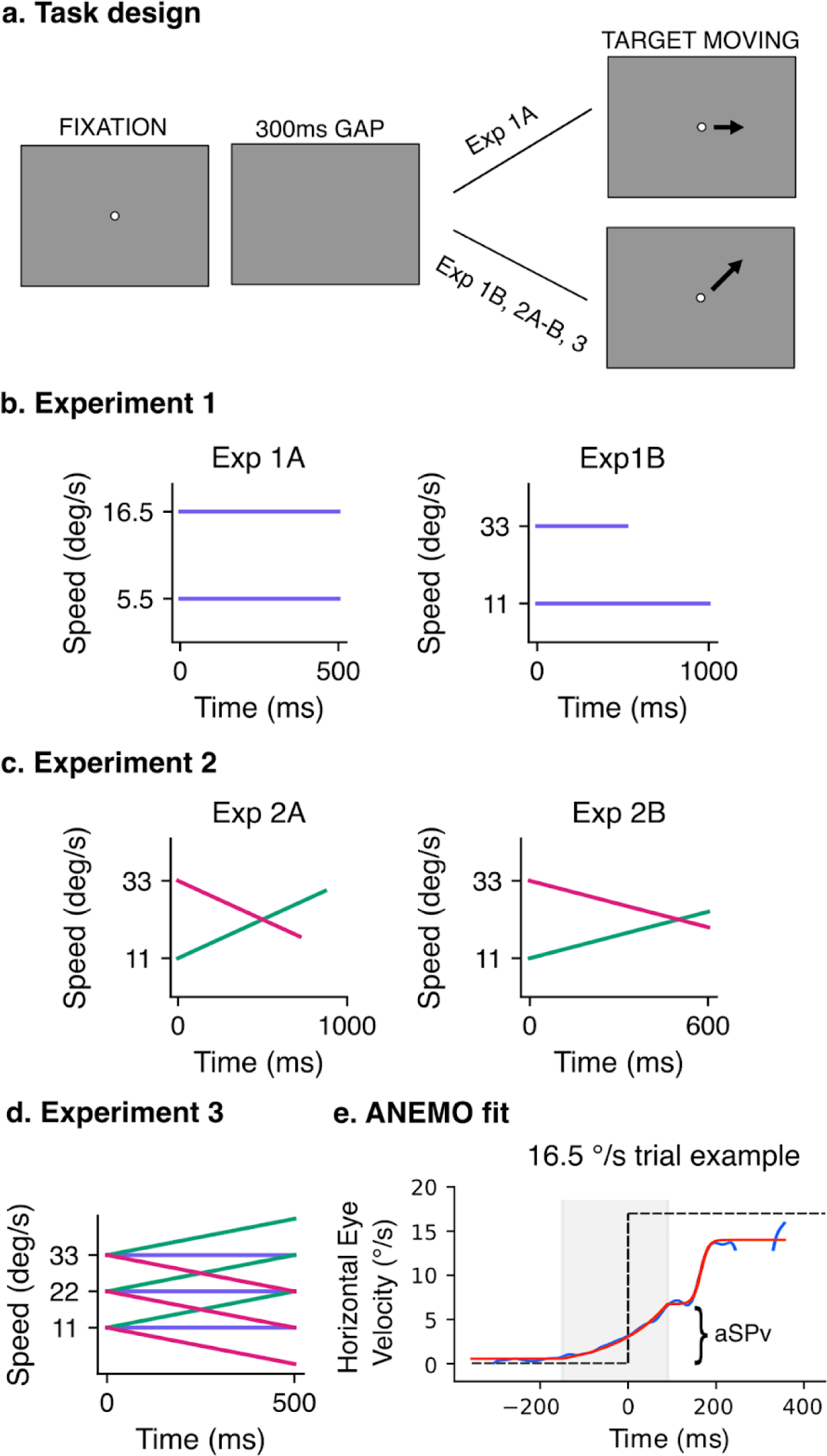
Experimental designs. **(a).** Each trial started with a fixation point displayed at the center of the screen for a random period, followed by a gap of 300 ms. The target then appeared at the center of the screen and started moving. **(b). Experiment 1.** In Experiment 1A, the target moved horizontally to the right at one of the two constant speeds (5.5 or 16.5 °/s, in blue). The probability of a high-speed (P(HS), v=16.5 °/s) vs a low-speed P(LS)=1-P(HS), v=5.5 °/s) trial was varied between experimental blocks with 7 different values of P(HS) (0, 0.1, 0.25, 0.5, 0.75, 0.9, 1). In Experiment 1B the target always moved in a fixed direction, chosen between one of the four diagonals (counterbalanced between participants). We displayed two different constant target speeds (v11c and v33c, of 11 and 33 °/s, respectively, in blue). Conditions were designed with a constant target displacement, and we manipulated the probability of the target kinematics in each experimental block with the probability of v11c spanning the values (0, 0.3, 0.7, 1) and P(v33c)=1-P(v11c). **(c) Experiment 2.** Experiment 2A followed a design similar to Exp 1B, but this time the target kinematics included acceleration conditions. The initial speed was kept the same as in Exp 1B, but the target with initial speed at 11°/s accelerated (v11a, a=22 °/s^2^, in green) and the target with initial speed at 33 °/s decelerated (v33d, a=-22°/s^2^, in pink). In Experiment 2B, target kinematics were the same as in Exp 2A, except the time duration of the target motion was kept constant at 600 ms. **(d)** Experiment 3. We displayed three different initial speeds (v11, v22, v33 of 11, 22 and 33 °/s, respectively) combined with three different acceleration values (a, c and d, of 22, 0 and −22 °/s^2^, in green, blue and pink respectively) in fully predictable blocks. Target motion duration was kept constant and lasted 500ms. **(e) Model fitting of eye velocity profiles in individual trials.** The bottom panel shows an example of the eye velocity trace (blue curve) and of the ANEMO model fit (red curve, Pasturel et al., 2018) in an individual trial. The selected model fitted an exponential function to the anticipatory phase, and a sigmoid to the initial visually-guided pursuit phase. From this model we extracted the maximum of the anticipatory velocity (aSPv). The model fitting procedure for Experiments 1B, 2A, 2B and 3 was similar to Experiment 1A.

#### Experiments 1B, 2 and 3

##### Experiment 1B-2A

The same group of participants took part in Experiments 1B and 2A. Stimuli were presented using the Psychtoolbox (Brainard, 1997) package for MATLAB. A Display++ monitor (CRS Ltd., Rochester, UK) with a refresh rate of 120 Hz was placed at 57 cm distance in front of the participant. Eye movements were recorded using an Eyelink1000, an infrared video-based eye tracker (SR Research Ltd., Ottawa, Canada). This experiment probed aSP with different target constant and varying speed conditions, while manipulating the probability of each condition. With respect to Experiment 1A, a larger pool of participants was tested with a different set of target speed values and a smaller set of speed probabilities and fewer trials per condition. Because several studies have reported anisotropies for saccadic and smooth pursuit eye movements across different directions (e.g., Grasse & Lisberger, 1992; Ke et al., 2013; Rottach et al., 1996; Takeichi et al., 2003), we decided to investigate whether the results observed in Experiment 1A can be generalized across target motion directions. Accordingly, in Experiment 1B-2A the target moved along one of the four diagonal directions, counterbalanced between participants.

**Figure 1a** shows the experimental design. Each trial started with a white fixation dot in the center of the screen for a random interval between 300 ms and 600 ms. This fixation period was followed by a 300 ms gap. At the end of this period, the target (a white circle of 0.6° diameter, maximum luminance 45 cd/m^2^ on a black background with luminance <1 cd/m^2^) appeared at the center of the screen and started moving in one of the four diagonal directions (always the same for one participant) with different target kinematics conditions: the target speed was either constant (v11c and v33c; 11 and 33 °/s respectively), uniformly accelerating (v11a; starting from 11 °/s, a=22 °/s^2^), or decelerating (v33d; starting from 33 °/s, a=-22 °/s^2^). In Experiments 1B and 2A, target motion duration was adapted to the target kinematic properties in order to achieve a similar spatial displacement on the screen across conditions: in practice target movement lasted 1 and 0.52 s for v11c, and v33c, respectively, 0.87 s for v11a, and 0.72 s for v33d.

Participants first completed 4 blocks of 100 trials with a single target kinematics condition (v11c, v33c, v11a, v33d, presented in randomized order across participants). Notice that these conditions can be considered as a probability of 1 for each kinematic value. Next, the probability of the different target kinematics was manipulated across blocks of 200 trials each. For blocks with constant speed, we used v11c and v33c, with P(v33c) being equal to either 0.3 or 0.7 and P(v11c)=1-P(v33c). The two probability levels were presented in random order across participants. For blocks with accelerating speed, P(v33d) could be either 0.3 or 0.7 (P(v11a)=1-P(v33d)). Again, these two probability conditions were randomly interleaved.

##### Experiment 2B

Having different motion durations across conditions might have introduced some confounds affecting anticipatory behavior. First, estimates of target acceleration can be impaired if the target motion is presented too briefly (Bennett, De Xivry, et al., 2010; Bennett et al., 2007). Second, if anticipatory behavior relies on the estimate of mean target velocity, rather than on its accelerating dynamics (Brouwer et al., 2002; Gottsdanker et al., 1961; Schmerler, 1976), then the duration of the visual motion epoch might influence the mean velocity estimate for targets with accelerating speed. Therefore, we ran an additional control experiment (Exp 2B) with the same design as Exp 2A but with one main difference: target motion duration was held constant (600ms), resulting in different target final positions across conditions.

##### Experiment 3

While Experiments 1 and 2 were designed to probe the effects of probability manipulation for both constant and accelerating target motion, we did not directly address whether and how the accelerating motion of predictable accelerating targets drive anticipatory eye movements. Most of the literature on smooth pursuit focus on constant speed targets, and the very few studies manipulating acceleration focuses on the effects of this manipulation on the visually-guided pursuit (Brostek et al., 2017; Krauzlis & Lisberger, 1994) or short-term predictive tracking (e.g. blanking paradigm, Bennett et al., 2007, 2010; Bennett & Barnes, 2006; Bennett & Benguigui, 2013). Interestingly, Bennett & Barnes (2006) showed that for a fixed initial target speed and when accelerating target motion is presented in a blocked design, anticipatory eye movements scale with target acceleration. In order to better understand the relationship between target acceleration and anticipatory eye movements, we ran Experiment 3 (**Figure 1d**), with a *fully-crossed* design, in which the target could take one out of three possible initial speeds (v11, v22, and v33 of 11, 22, and 33 °/s, respectively) and one out of three possible acceleration values (of 22, 0 and −22 °/s^2^, labeled as “a” for *accelerating*, “c” for *constant* and “d” for *decelerating,* respectively). This design yielded in nine blocks of fully-predictable target motion. In all blocks, the target motion duration was held constant to 500 ms. Participants completed 100 trials in each block with v11 and v22 as initial speeds, and 120 trials in each block with v33 as initial speed. The small difference in the number of trials was introduced to compensate for the higher number of excluded trials for the data analysis, due to a higher occurrence of saccades around target motion onset, a phenomenon already observed in the previous experiments when the fastest speed (v33) was presented.

### Eye-movements recording and preprocessing

For Experiment 1A, the analog voltage measure collected with the scleral coil technique and reflecting the right-eye rotation was low-pass filtered (DC-130 Hz) and digitized with 16-bit resolution at 1000 samples per second (to obtain the eye’s horizontal and vertical position). For Experiments 1B, 2A, 2B and 3, the right-eye horizontal and vertical position was recorded with an infrared video-based eye tracker, EyeLink 1000 (SR Research), at 1 kHz.

For all sets of recorded eye movements, position data was converted in an ASCII format. After conversion, the ANEMO toolbox (Pasturel et al., 2018) and custom-made python scripts were used to pre-process the data. Position data was low-pass filtered (acausal second-order Butterworth low-pass filters, 30 Hz cut-off), numerically differentiated to get the eye velocity in degrees per second and then de-saccaded using ANEMO’s implementations. In practice, saccade detection was implemented by jointly applying the absolute eye velocity threshold criterion (30°/s, by default in the Eyelink system) and the relative velocity threshold (akin to the method proposed by Engbert & Kliegl, 2003). The epochs corresponding to detected saccades were removed from analysis (given not-a-number values). Trials with more than 40% of missing data points in the [-100,200] ms window around the target onset for Exp 1A and more than 70% of missing data points in the [-100,100] ms window around the target onset for Exp 1B, 2A,B and 3, or with more than 60% of missing data points overall (for all experiments), were automatically excluded from the pre-processing pipeline. ANEMO was used to fit a piecewise linear and non-linear model to individual trials’ eye velocity to extract the relevant oculomotor parameters (see **Figure 1e** for an illustration of the model fitting and extracted parameters). The model comprised a linear baseline phase (flat linear regression), and two non-linear phases: an anticipatory phase modeled by an exponential function and a visually-guided phase modeled by a sigmoid function. Importantly, we selected this model after comparing its performance quantitatively (using AIC, BIC and RMSE indicators of goodness of fit) with a more standard piecewise linear model (implemented in ANEMO): in all experiments our model (linear + non-linear model) performed better than the linear one in the large majority of trials. In Experiment 1A, the fit was performed in the time window of −300 to 350 ms relatively to the target motion onset. In Experiments 1B, 2A,B and 3, the fit was performed in the time window of −300 to 300 ms relatively to the target motion onset. The relevant model-fit parameter for this study was only the maximum velocity of the anticipatory phase (aSPv), calculated as the maximum value of the exponential function (i.e. its value at the offset of the anticipatory phase). Other oculomotor parameters are automatically extracted by the ANEMO toolbox, but we do not report them in the present study.

Eye velocity traces and model fits were visually inspected to exclude the remaining aberrant trials and those with extremely poor fits. Overall, 12% of the trials were excluded on average for Experiment 1A (median: 10%, max: 18%), 16% for Experiment 1B-2A (median: 14 %, max: 32%), 13 % for Experiment 2B (median: 13 %, max: 15 %), and 26 % for Experiment 3 (median: 25 %, max: 33 %).

### Data analysis

In all experiments, when analyzing the effects of the probability bias, the first 10 trials of each block were excluded from analysis. This was done in the aim of excluding large fluctuations related to learning the stimuli statistics and focus on average values rather than on their rate of change. For control, we also repeated all analyses excluding the first 50 trials: the results did not change significantly.

Linear Mixed Effects regression Models (*nlme* package for R) were used to evaluate the effects of the target kinematics (different constant speed and acceleration conditions) and of the kinematic probability on the anticipatory smooth pursuit velocity (aSPv) estimates. Because not including a true random effect can increase the false positive rate (Barr et al., 2013), participants were treated as a random effect and all fixed effects were allowed to vary with it. Since this approach usually leads to models that don’t converge because of the high number of parameters and the correlations between them, when needed, we used the *buildmer* package for R (Voeten, 2020) to find the maximal model (i.e., the model including the most of variables) which still converges for the dependent variable. After finding the models, we fitted the data and result tables were exported with the *stargazer* package for R (Hlavac, 2022).

For Experiment 1A, the models included only speed probability as a fixed effect for the oculomotor anticipatory velocity (aSPv). We chose to use the probability of the highest speed, P(HS), as the independent variable in the model 1:

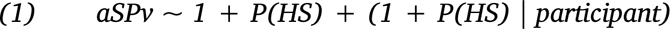

The variable before the ∼ symbol is the dependent variable, and the variables after it are the independent variables (also called fixed effects). The 1 corresponds to the model intercept. For the variables within the parentheses, each one before the | symbol is allowed to vary for each level of the variable after it (also called random effect). In other words, for each participant, the model will return a different best-fit value for the intercept and the slope of the linear dependence upon probability. For the analysis of the probability-mixtures of Experiment 1B % (v11c vs v33c) and Experiment 2 (v11a vs v33d), we also added the axis (horizontal/vertical) as an interaction factor with the probability, given that the target moved along one of the diagonals. For Experiment 2, we included an interaction with the experiment (2A/2B). The final models, as well as the statistics tables are presented in the supplementary material.

It is known that recent trial-history can affect the behavior in the present trial (sequential effect), and this could be seen as a confound in our analyses addressing the effect of the target motion probability across several trials. Therefore, we re-ran the regression models for Experiments 1A and 1B adding the target velocity at trial N-1 (*Tv_N-1_*) as an interaction variable:

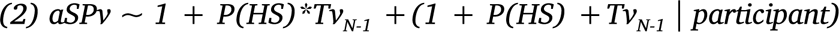

In Experiment 3, considering that each target kinematic condition could be uniquely identified as a combination of an initial velocity and acceleration values, we ran a second linear mixed-effects regression model: this time the main independent variables were the target initial velocity (*v0*: 11, 22, or 33°/s) and the target acceleration (*accel*: 0,+22 and −22°/s^2^), as well as their interaction, all treated as parametric variables:

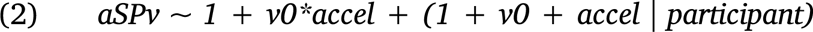

In order to test the actual role of target acceleration this model was tested against the model including only v0 as fixed effect using the *bayestestR* package for R (Makowski et al., 2019). Pairwise comparisons between aSPv corresponding to the target kinematics (*Tk*) conditions of Experiment 3 were based on a categorical LMM, initially defined as:

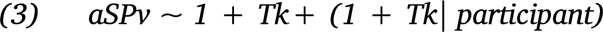

Contrasts between the different conditions were performed using the *emmeans* package for R (Lenth, 2017) and p-values were adjusted for multiple comparisons using the Benjamini & Hochberg method for controlling the false discovery rate.

We then tested the hypothesis that anticipatory eye velocity is driven by an estimate of the mean target velocity across a finite *temporal window of integration* (TWI) starting at target motion onset (time 0) and ending at time TWI^end^. In order to do so we fitted, for the pooled participants’ data, the following linear regression:

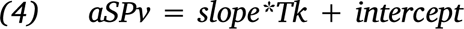

and extracted the individual best-fit value for the parameters *slope_i_* and *intercept_i_*. Here, Tk included v11c, v22c, and v33c. We then used the same relation (5) for the six conditions with accelerating targets, indexed by the suffix i, replacing Tk by the target speed estimate (TSE***_i_***), which approximates the accelerating kinematics:

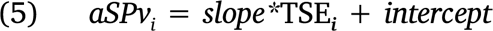

Solving with respect to TSE***_i_*** we have:

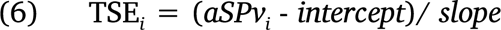

By imposing the equality between the estimated target speed TSE***_i_*** and the mean of the accelerating/decelerating target speed over a finite TWI, between 0 and TWI^end^, we obtain:

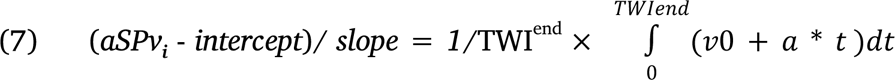

where a is the acceleration value and v0 the initial target speed for accelerating conditions. Finally, by solving Equation 8 with respect to TWI^end^ we could estimate, for each acceleration condition, the final point in time of the temporal window of integration,

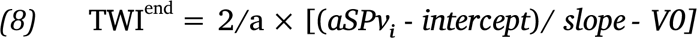

In order to have a robust estimate of the variability of such temporal window of integration across the population, we used the bootstrapping technique (n=1000, Efron, 1979) and extracted the 95% confidence interval around the mean estimated TWI^end^.

## Results

### Effects of target speed probability on anticipatory Smooth Eye Movements

Using a state-of-the-art eye movement recording technique (scleral search coil), we first investigated the effects of target speed probability upon anticipatory smooth eye movements. In Experiment 1A, three participants had to smoothly pursue a target which moved rightwards along the horizontal axis with two different possible speeds (referred to as high-speed, HS, and low-speed, LS) randomly interleaved across trials but with a given probability of occurrence (P(HS) and P(LS)=1-P(HS)) within a block. **Figure 2a** shows the trial-averaged eye velocity curves for one participant, sorted by probability and target speed conditions. Each probability condition is represented by a different color. The two different target speed profiles are illustrated by the horizontal dotted lines; time zero corresponds to target motion onset. Participants were able to track the two target motions with high accuracy, as shown by the convergence of eye velocity to the target speed during steady-state pursuit. Since target motion direction was always rightward, we observed a strong anticipatory response for all speed probability conditions, as evidenced by the non-zero eye velocity at the usual pursuit latency in humans (∼100 ms). However these anticipatory responses in the predicted direction were also modulated by target speed probability. We analyzed such modulation by considering the relationship between amplitude of anticipatory responses and P(HS), i.e., the probability of the highest speed (16.5 °/s). As illustrated in **Figure 2a**, higher values of P(HS) drove stronger anticipatory pursuit, regardless of the actual target speed (16.5 ou 5.5 °/s). **Figure 2b** plots the relationship between mean anticipatory eye velocity (aSPv) and P(HS), for the 3 participants. The gray curves are the linear relationships estimated from the Linear Mixed Effects Model (LMM). We found a clear linear dependency of anticipatory response upon the probability of target speed, in the direction of target motion. The LMM statistical analysis (with P(HS) as a fixed effect) shows that aSPv significantly increased with higher probability (P(HS) effect: beta = 3.48, 95% CI = (3.14,3.84), p<.001). Overall, anticipatory responses were stronger by ∼200 %, rising from 2.5 to 7.5 °/s when P(HS) increased from 0 to 1.

**Figure 2.**
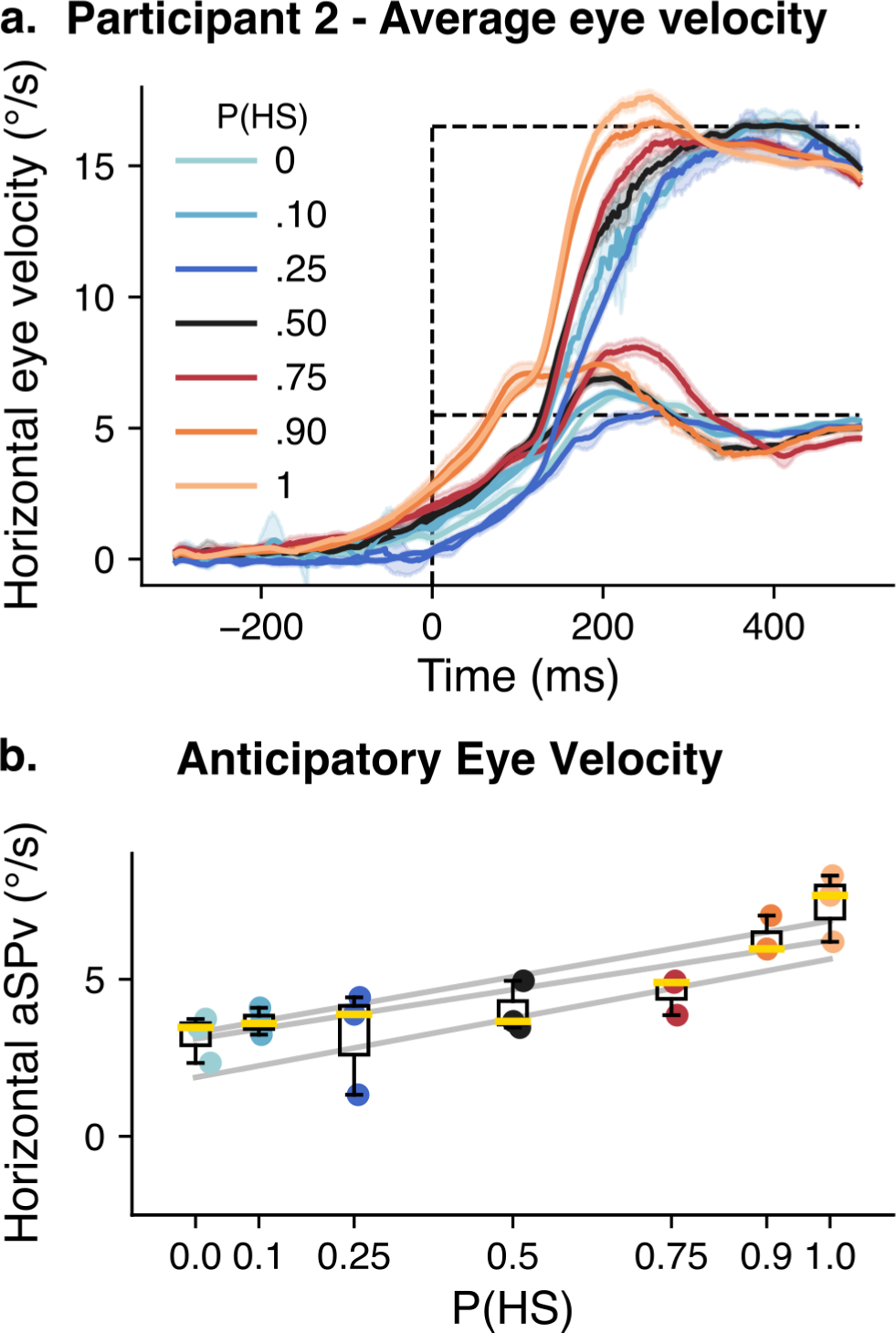
Experiment 1A: Dependence of anticipatory eye velocity upon target speed probability. **(a)**. Average eye velocity as a function of time (+/− 95% confidence interval) for one representative participant. Trials are grouped by probability of the higher speed (HS). Each color corresponds to one probability condition. The time zero corresponds to the target onset, and the dashed lines indicate the two possible target speeds. **(b)**. Anticipatory eye velocity for the group of participants. Each box plot (median in yellow, box limits corresponding to the 25% and 75% quartiles, and whiskers corresponding to 1.5 times the interquartile range, IQR) corresponds to one probability condition. The gray lines show the linear mixed model fit for each participant.

In a second version of this Experiment (Exp 1B), our objective was to replicate the speed-probability dependency of anticipatory pursuit on a larger group of 13 participants, with a less invasive eye tracking technique. We used two new speeds (11 and 33°/s) and a reduced set of probability conditions in the mixture blocks (P(v33c)=0, 0.3, 0.7 and 1). By using an oblique target motion direction, our second objective was to generalize this speed probability dependency across visual motion directions, by having the target moving along one of the 4 diagonal directions (**Figure 1b**). To directly compare the results of Experiment 1A and 1B, we first report the effects of target speed probability observed when varying P(v33c) – and therefore P(v11c)=1-P(v33c) – across blocks with constant speed (v11c,v33c) mixtures. **Figure 3a** shows an example of the trial-averaged eye velocity as a function of time for one participant, sorted according to the target velocity profiles (dotted lines) and P(v33c) values. Horizontal dotted lines depict the target horizontal and vertical velocity components (i.e. 7.77 and 23.3 °/s for 11 and 33 °/s radial speed). A comparison with **Figure 2a** shows a behavior similar to Exp 1A where target motion was horizontal. Final pursuit velocity matches target velocity, especially for the lowest target speed (v11c). The fact that steady-state eye velocity gain remained lower than 1 for the fastest target speed (v33c) is consistent with previous studies (Carl & Gellman, 1987; Dodge, 1930). As in Exp 1A, target motion direction remained constant within a block, thus a robust anticipatory pursuit response was always observed. However, its amplitude increased when P(v33c) increased. Such dependency is illustrated in **Figure 3b**, where horizontal and vertical components of anticipatory eye velocity (aSPv) are plotted against P(v33c). Both components increased linearly with the probability of the highest speed. A symmetric relationship was observed with P(v11c). We ran the LMM statistical models for the anticipatory response (aSPv), including the effect of P(v33c), as for Exp 1A. We added the effect of the eye velocity axis (horizontal or vertical) and its interaction to test whether aSPv was differently modulated along the horizontal and vertical dimensions. aSPv increased significantly for higher probability of P(v33c) (**Figure 3b**, P(v33c) effect: beta = 2.74, 95% CI = (1.82, 3.66), p<.001). We did not find a significant difference (given a criterion of alpha <0.01) between axes (main axis effect: beta = −0.74, 95% CI = (−1.46, −0.016), p=0.046), but the effect of P(v33c) was significantly smaller in the vertical axis (P(v33c)*axis effect: beta = −0.67, 95% CI = (−0.98, −0.36), p<.0001). Overall, the two experiments strongly support the fact that anticipatory eye velocity scales with the probability of target speed, in a similar way across different motion directions in the plane, although slightly less robustly along the vertical direction.

**Figure 3.**
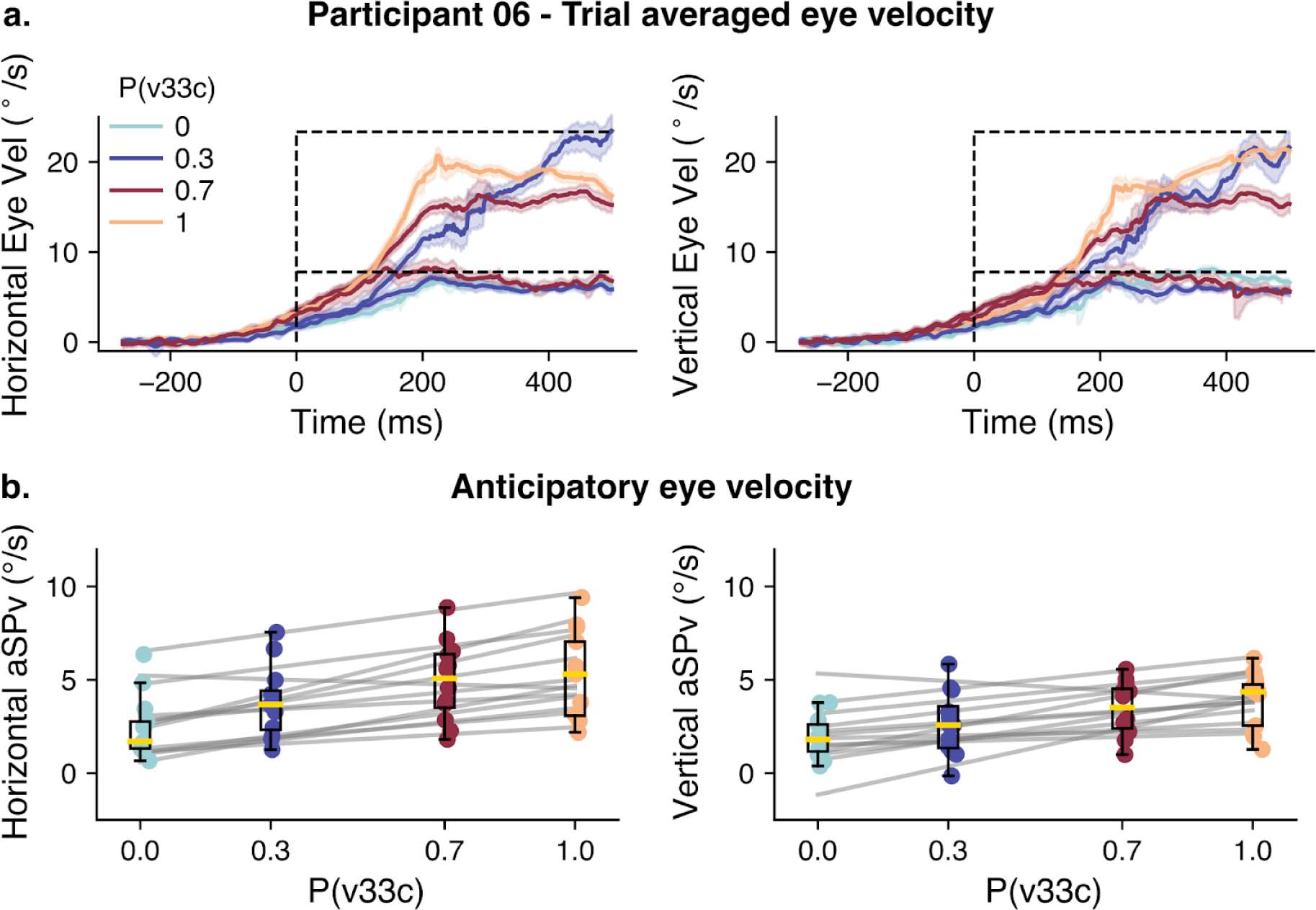
Experiment 1B: Dependence of anticipatory eye velocity upon target constant-speed probability. **(a).** Average eye velocity for trials grouped by probability of v33c and by target velocity for the (v11c,v33c) mix. The left panel shows the horizontal eye velocity, while the right panel shows the vertical eye velocity for a representative participant. Each color corresponds to the different probabilities of v33c. **(b).** Amplitude of anticipatory pursuit is plotted against P(v33c), along the horizontal (left panel) and vertical (right panel) axes. Data represented in the same way as Figure 2.

#### Short and long-time scale factors affecting speed expectation and eye movement anticipation

Recent trial history, that is the stimulus properties (e.g. the target speed) observed in the previous trial, or across the few previous trials, can modulate anticipatory eye movements (e.g. Heinen et al., 2005; Kowler & McKee, 1987; Kowler & Steinman, 1979; Maus et al., 2015). Importantly, several studies have also shown that both short-term factors related to one or few previous trials, and longer-term factors, related to global statistical estimates can coexist and interact to control perception and visuomotor behavior (e.g. Chopin & Mamassian, 2012; Maus et al., 2015, Damasse et al., 2018; Falmagne et al., 1975; Kowler, 1984; Pasturel et al., 2020; Wu et al., 2021). In order to quantify the effects of the previous trial’s speed on the anticipatory eye velocity in the present study, as well as its interaction with the block’s speed probability, we ran a new LMM, now including both the speed-probability and the target speed at the trial N-1 (Tv_N-1_). Note that our study was not designed specifically to study sequential effects (i.e., by presenting all possible combinations of N-1, N-2, N-3,… trials), and therefore we limit our analysis of short-term effects only to the effect of the trial N-1. When the previous trial was a low-speed trial, aSPv decreased when compared to a previous high-speed trial (Exp1A: beta = −0.50, 95% CI = (−0.66, −0.34), p<.001; Exp1B: beta = −0.61, 95% CI = (−0.88, −0.34), p<.001). For both Experiment 1A and 1B, we found that the main effect of target speed probability upon aSPv remained significant (Exp1A, Tv_N-1_=HS: beta = 4.35, 95% CI = (3.98, 4.72), p<.001; Exp1B, Tv_N-1_=v33: beta = 1.38, 95% CI = (0.44, 2.32), p<0.01). The interaction between the previous trial’s speed and the block’s speed probability was also significant, although with a different sign: for Experiment 1A the probability effect was reduced when the previous trial was low-speed compared to high speed (beta = −2.74, 95% CI = (−3.10, −2.38), p<.001); in contrast, for Experiment 1B the probability effect increased when the previous trial was at v11c compared to v33c (beta = 0.82, 95% CI = (0.38, 1.26), p<.001). These results are illustrated in the left panels of **Figure 4a,b**. In the right panels of **Figure 4a** and **4b** we plotted, for Experiment 1A and 1B respectively, the difference between the mean aSPv for trials following a high-speed (in red) or a low-speed (in blue) trials and the mean aSPv across all trials in a probability block. This illustration allows us to immediately capture how the N-1 trial’s effect is modulated across the probability values: a high-speed previous trial has a larger excitatory impact on subsequent anticipatory velocity when high-speed trials are less frequent. The symmetric interaction is observed for a low-speed previous trial, namely its inhibitory effect is stronger in blocks with a low probability of low-speed trials.

**Figure 4.**
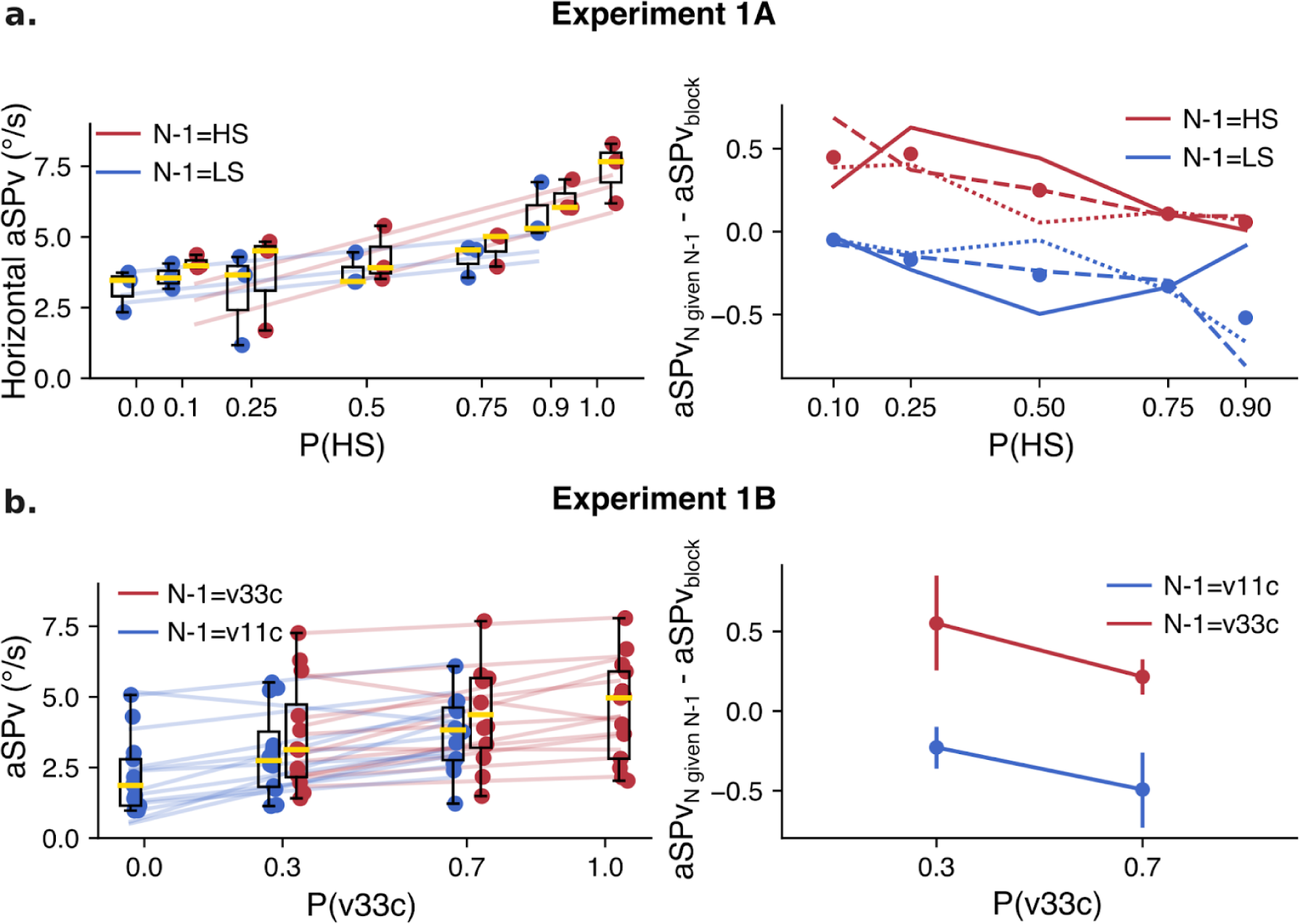
Effect of previous trial’s speed versus block’s probability. **(a) Experiment 1A.** The left panel shows the probability effect on the aSPv given that the trial N-1 was a high-speed trial (HS, red) or a low-speed trial (LS, blue). The right panel shows the difference between the mean aSPv for trials with N-1=HS (red), or N-1=LS (blue) and the mean aSPv in the whole block. Each trace corresponds to one participant, while dots correspond to the average between participants. **(b)** Same analysis for Exp 1B. On the right panel, individual traces were substituted by the 95% confidence interval across participants for the sake of clarity.

### Effects of the accelerating target probability on anticipatory eye movements

Following the same reasoning as for constant speed mixture blocks, we tested the effect of the probability of accelerating target kinematics on aSP. In Experiments 2A and 2B, we compared different probabilistic mixtures of trials of accelerating or decelerating target motion. Participants ran blocks of 4 different probability pairs (P(v33d), P(v11a)): (0,1), (0.3, 0.7), (0.7, 0.3) and (1,0). In Figure 5, we present the results relative to P(v33d) values. Figure 5a shows horizontal and vertical trial-averaged eye velocities for one participant and each available combination of P(v33d) and target kinematic conditions. Again, each color depicts one probability condition and time zero indicates the target movement onset. We can see clear anticipatory responses, with stronger anticipation occurring for higher probabilities of the highest initial velocity and decelerating motion. After the anticipatory phase, eye velocity traces corresponding to vacc or vdec trials separate and converge to the target’s velocity which keeps changing in time according to the acceleration condition. Notice that the anticipation seen with P(v33d)=0 (i.e., P(v11a)=1) was particularly small but still significant in participant 6, as in all others. Figure 5b plots the horizontal and vertical aSPv, as a function of P(v33d), for all participants. There is an increase in the amplitude of anticipatory pursuit as P(v33d) increases, as confirmed by the LMM statistical analysis (P(v33d): beta = 1.88, 95% CI = (1.11, 2.65), p<.001). Consistently with the previous analysis, we did not find any significant difference between axes (axis main effect: beta = −0.46, 95% CI = (−1.26, 0.34), p=0.26) and the interaction between axis and probability did not survive the model selection procedure, suggesting it was not significant. We found that the aSPv was slightly higher in Exp 2B (constant target duration) than in Exp 2A (constant target displacement) (experiment effect: beta = 1.68, 95% CI = (1.41, 1.95), p<.001), but, again, the interaction between experiment and probability was excluded from the selected model. Overall, in the probability-mixture blocks with accelerating targets, we observed a robust probability-dependent anticipation, similar to the mixture blocks with constant speed and this regardless of the motion direction. The significant main effect of the experimental design (difference between Exp 2A and 2B) suggests that the temporal regularity across trials generally favors anticipatory behavior; however, the lack of interaction between the experiment design and the probability effect argues against a critical role of the motion presentation duration (at least in the tested range, namely above 500ms) on the integration of information about the target acceleration.

**Figure 5.**
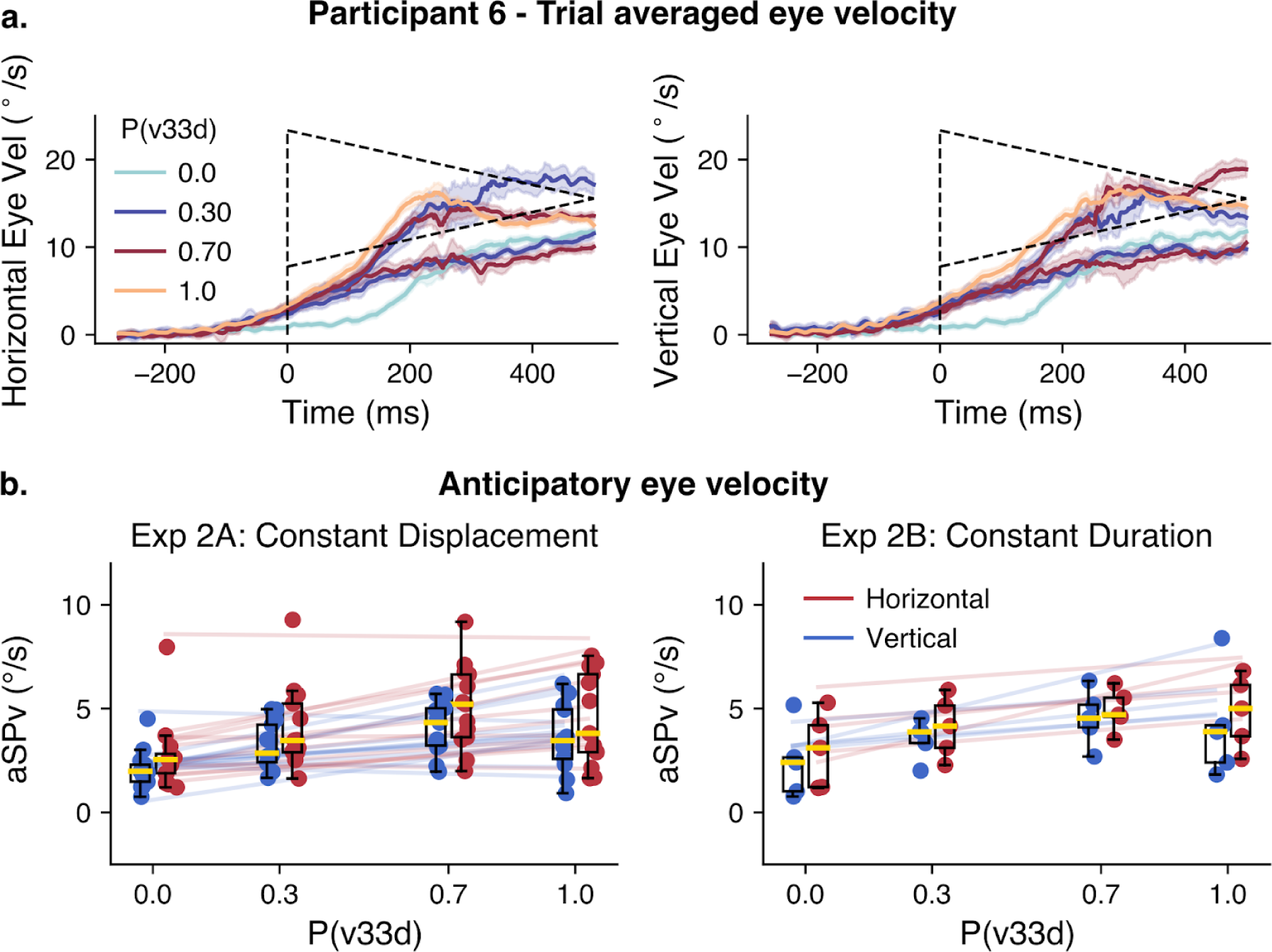
Experiment 2A,B. Effect of the probability of accelerating target kinematics upon anticipatory eye movements (mixture blocks). **(a).** Average eye velocity across time for a representative participant. Trials are grouped according to the probability of v33d (P(v33d)) and sorted by the different target kinematic conditions for each (P(v33d), P(v11a)) mixture. Left and and right panels show horizontal and vertical eye velocity profiles, respectively. The dashed lines show the target velocity. **(b).** Mean anticipatory eye velocity as a function of the probability of v33d, group results. The left panel shows the data for Exp 2A, while the right panel shows the data for Exp 2B. Horizontal data is shown in red, and vertical data is shown in blue. Data represented in the same way as Figure 2.

### Effects of different target kinematics on anticipatory Smooth Eye Movements

Lastly, we questioned in Experiment 3 how the anticipation for accelerating target motion compares to that observed for constant target speeds. We compared three conditions where oblique target motion had a constant speed (v11c, v22c and v33c, corresponding to radial 11, 22 and 33 °/s, respectively) to conditions in which the target started at 11, 22 or 33 °/s and either accelerated uniformly (v11a, v22a, v33a, acceleration = 22 °/s^2^) or decelerated uniformly (v11d, v22d, v33d, acceleration = −22 °/s^2^). The different target kinematic conditions were presented in a block design and motion direction was fixed for each participant, leading to full predictability of both target’s trajectory and kinematics (P=1). **Figure 6** shows eye velocity profiles recorded in one participant, for Exp 3, for each target kinematic condition illustrated by the dotted lines. Each row of **Figure 6** corresponds to one initial target speed value, while acceleration values are shown in different colors (accelerating motion in green, constant motion in blue, and decelerating motion in pink). Predictable constant speed targets drove strong anticipatory pursuits that were scaled according to target speed (blue curves, from top to bottom). Moreover, accelerating conditions also resulted in clear anticipatory pursuit responses. As clearly seen with the 22 and 33°/s initial speed conditions, green and red curves are, respectively, above and below the blue ones, illustrating that anticipatory responses were modulated by both initial speed and target acceleration.

**Figure 6.**
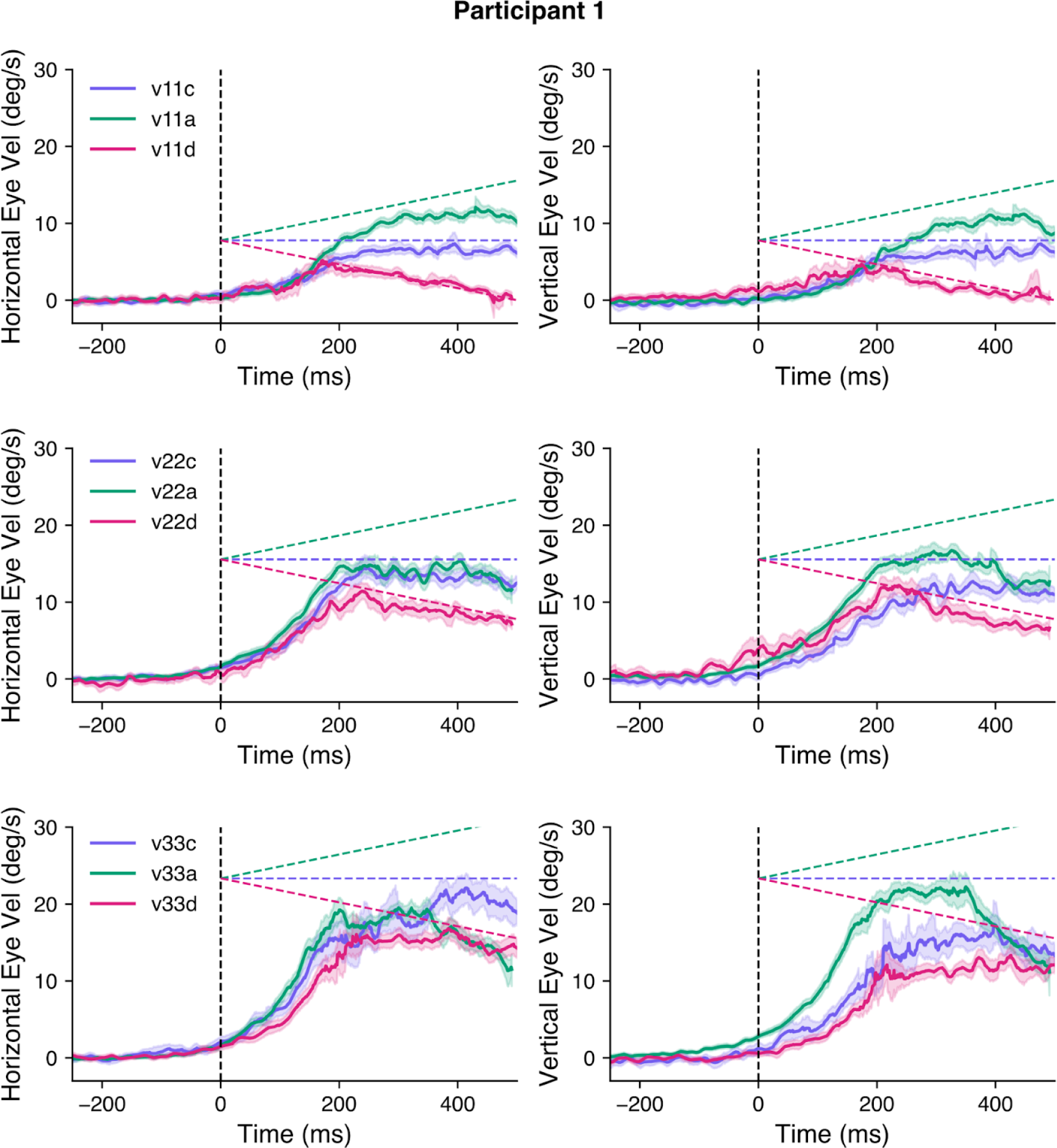
Experiment 3: Effect of target kinematics on aSP (fully predictable blocks). **(a).** Average eye velocity over time with trials grouped by target speed (rows: v11, v22, and v33 from top to bottom) and target acceleration values (color coded: accelerating in green, constant in blue, decelerating in pink) for one participant of Exp 3. The dashed lines show the corresponding target speed profile for each acceleration value.

**Figure 7** illustrates the amplitude of anticipatory pursuit responses (aSPv) for the different conditions across all participants. Note that, for each initial speed, there is a tendency for aSPv to increase as the acceleration increases. To statistically test the effect of acceleration on the anticipatory eye velocity, we ran a parametric linear mixed-effects regression model including both the acceleration and the initial target speed as independent variables.

**Figure 7.**
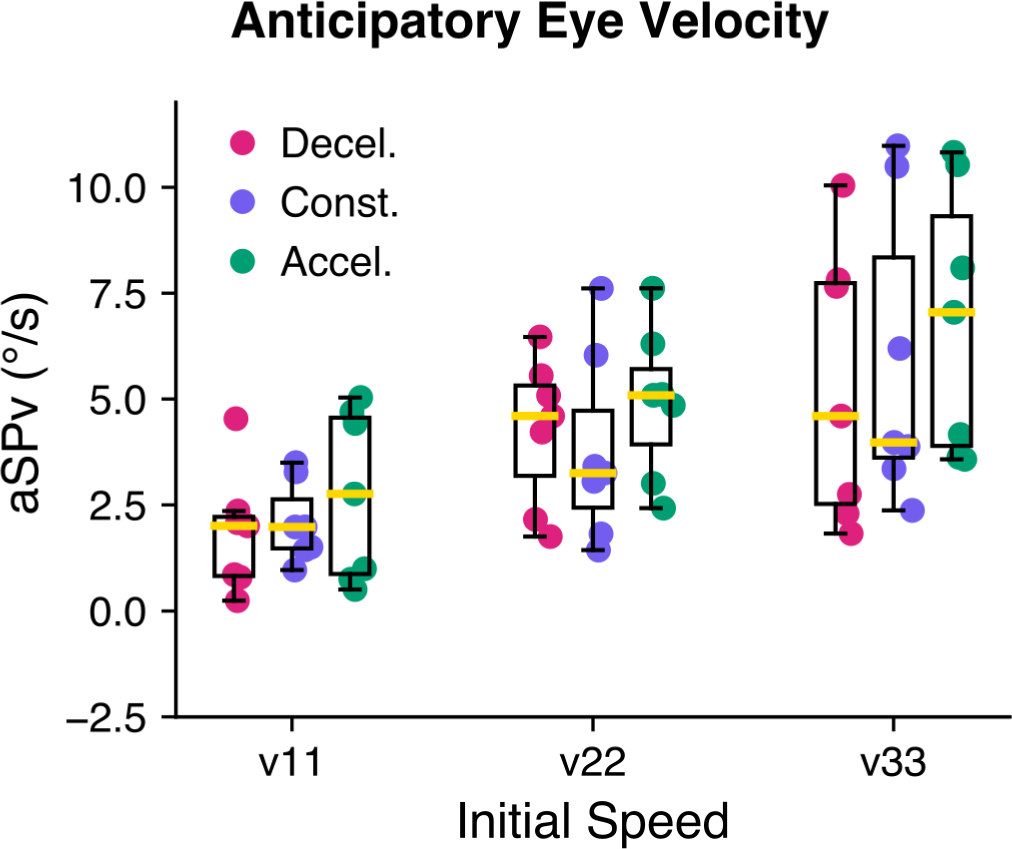
Experiment 3: Group effect of target kinematics on the aSPv. aSPv is grouped by initial velocity (v11, v22 and v33, from left to right, respectively) and acceleration values are shown in different colors (decelerating target motion in pink, constant target motion in blue and accelerating target motion in green). Data are represented in the same way as Figure 2.

We found that a model including both the initial speed and acceleration was significantly better than a model including only the initial speed (speed-only model BIC = 43521.6; full model BIC = 43308.97; Bayes Factor = 1.48e+46). The initial speed significantly modulated the aSPv (v0: beta = 0.24, 95% CI = (0.15, 0.34), p<.001), while the acceleration alone had a significant but smaller positive effect on the aSPv (accel: beta = 0.017, 95% CI = (0.004, 0.030), p=0.009), indicating that indeed aSPv increases with target acceleration. We also found a significant positive interaction between initial speed and acceleration (v0*accel: beta = 0.001, 95% CI = (0.0004, 0.002), p=.004), indicating that the effect of acceleration is stronger for high initial speeds when compared to low initial speeds.

#### Temporal window of estimation of the mean target speed for accelerating motion

Previous studies investigating predictive smooth pursuit during the transient disappearance of the target have compared different possible schemes of temporal integration to build an internal model of complex target motion. Such alternative internal models could either take into account only the last sample of observed target velocity before target blanking, or an average of velocity across a finite time window, or use both the last velocity sample and its rate of change. Their results suggest that the rate of speed change (i.e. acceleration) was only taken into account if target displacement properties were estimated during a sufficiently long interval (Bennet et al. 2007). Performance was in any case inaccurate due to a lack of sufficient extrapolation of accelerating motion (Bennet & Benguigui 2013). Together with the results of Bennet and Barnes (2006), our results suggest that the target’s acceleration is integrated, though with a small weight and large variability, in the internal model of visual motion that drives anticipatory smooth eye movements across a block of trials where the target motion is highly predictable (as illustrated in the previous section). However, such an effect does not imply, *per se*, that a representation of target acceleration is accessible to the visuo-motor system. For instance, an estimate of target speed across a finite temporal window could be used as a proxy for target motion and drive anticipation. Thus, we simulated a similar but more realistic version of the simplest, initial speed based, internal model of target motion. We reasoned that, rather than an instantaneous estimate of target velocity (in our case the target velocity at time 0, or target motion onset) the internal model of motion stored in memory would take into account an estimate of the mean target velocity computed over a finite time-window around target motion initiation. This alternative model could accommodate the observation that expectancy-based anticipatory velocity differs, but only weakly, for two targets with the same initial velocity and a different acceleration. Note that we can already speculate that such hypothetical temporal window of integration should be shorter than 500ms. If this was not the case, aSPv should not differ between target kinematic conditions that lead to the same mean target speed across the 500ms window of motion presentation. In Experiment 3, two pairs of conditions fit to this requirement, namely (v11a, v22d) and (v22a, v33d). For both pairs, aSPv is significantly higher for the second than for the first target kinematic condition (v11a-v22d = −1.503, SE = 0.139, p<.0001; v22a-v33d = −0.443, SE = 0.132, p<.001).

**Figure 8** illustrates the rationale and the results of our model-based analysis of the temporal window of velocity integration. We have shown in the previous sections that a linear regression describes the relationship between target speed and anticipatory eye velocity for predictable, constant target speeds (schematically illustrated in **Figure 8a**, upper panel, blue line). We assumed that 1) the same linear regression applies to the accelerating conditions as well, to describe the relationship between anticipatory eye velocity and the target speed estimate (TSE) approximating the accelerating kinematics; 2) such an estimate, TSE, would correspond to the mean target speed computed over a finite temporal window of integration (TWI), starting at time 0 (target motion onset) and ending at time TWI^end^. Thus, in order to have a reliable estimate of TWI^end^ for all kinematic conditions across participants, we first estimated the linear regression between aSPv and target speed at the group level, by pooling the data of all participants together. We then computed the mean TSE knowing the aSPv values corresponding to accelerating conditions and inverting the above-mentioned linear relation (as illustrated in the top panel of **Figure 8a**). From the TSE value, we finally inferred the mean TWI^end^ (as schematized in **Figure 8a**, bottom panel) and its variability using a bootstrapping procedure (Efron, 1979, see **Figure 8b**; for details of the calculations please refer to the figure caption and the Methods section). Note that the estimated TWI^end^ for accelerating conditions is between 0 and 500 ms, although the distribution is very broad (v11a, mean = 0.15, 95% CI = (−0.04, 0.34); v22a, mean = 0.18, 95% CI = (−0.03, 0.33); v33a, mean = 0.19, 95% CI = (−0.14, 0.49)). Remarkably, the TWI^end^ distribution for decelerating conditions displays smaller values, largely overlapping with 0, but also includes a wide range of unrealistic (negative) values (v11d, mean = −0.01, 95% CI = (−0.17, 0.10); v22d, mean = −0.09, 95% CI = (−0.23, 0.07); v33d, mean = 0.004, 95% CI = (−0.33, 0.32)).

**Figure 8.**
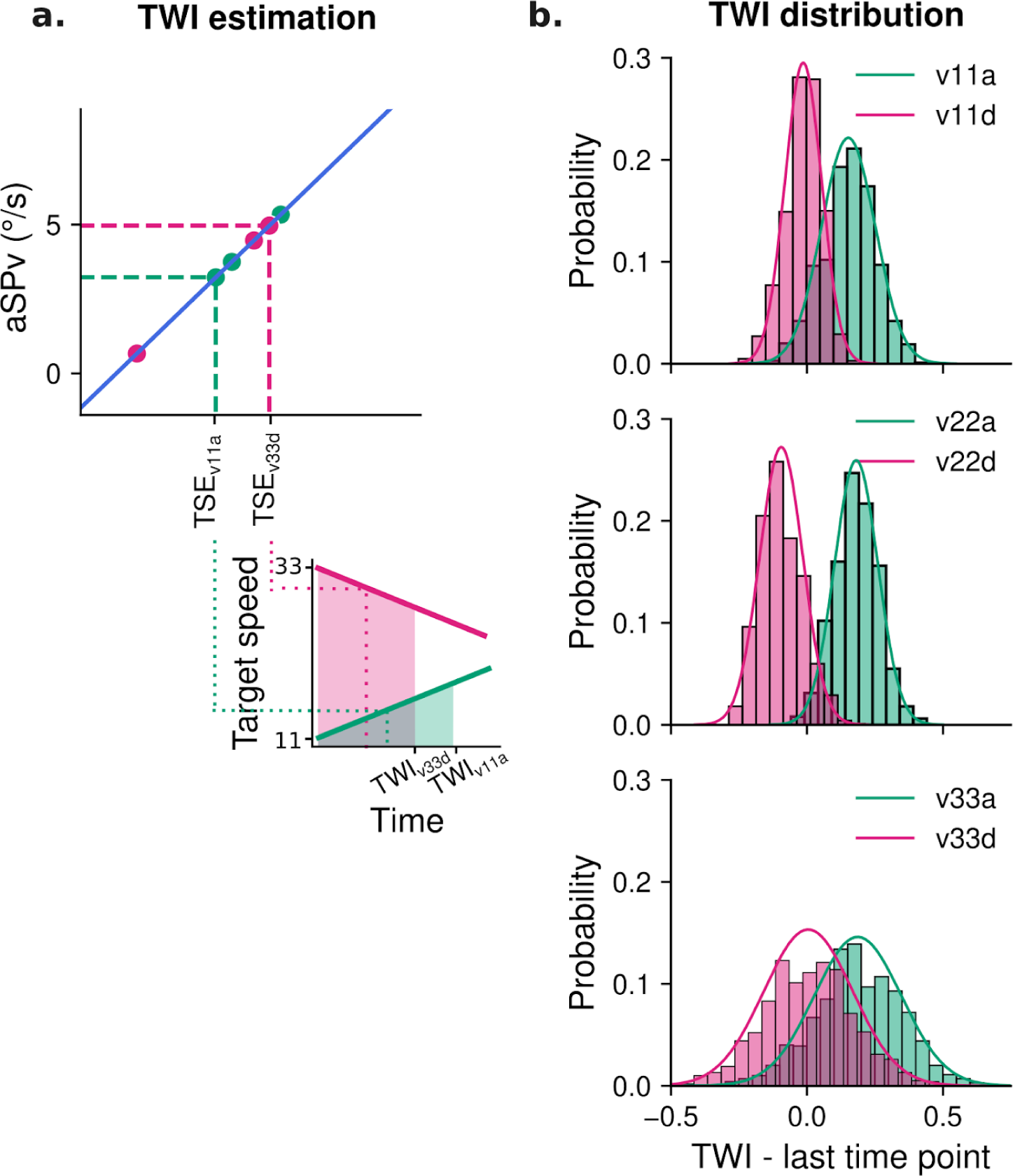
Testing the hypothesis of a finite temporal window of integration (TWI) to extract a target speed estimate (TSE) for accelerating conditions. (**a**) Schematic illustration of the procedure to compute the temporal window (the number of datapoints is reduced, for the sake of clarity: for details refer to the Methods section). **Upper panel**: At the group level and for the constant speed blocks, we calculated the linear regression between aSPv and target speed (blue straight line). On the basis of this regression, for accelerating blocks (pink or green dots), we performed the inverse operation to calculate the estimated target speed (TSE) that would have elicited the observed aSPv. **Lower panel**: The TWI is estimated as the temporal window over which the mean of the accelerating target speed equals TSE, as exemplified for two particular accelerating conditions (v11a and v33d). **(b)** Distribution of the bootstrapped estimates of TWI for the accelerating conditions. Each panel shows the TWI distributions for one initial speed (v11, v22 and v33 from top to bottom) and for the accelerating (pink) vs decelerating (green) target motion. The smooth line depicts a gaussian fit of the histograms.

Overall, these results cannot exclude the possibility that visual accelerating motion is approximated by the mean target speed estimate across a finite time window. However, the large variability of the inferred TWI and the presence of incoherent results, like negative TWI^end^ (especially for deceleration targets) impose a strong caution in the interpretation. We will further discuss this finding in the Discussion section.

## Discussion

We had two main objectives with the present study. First, we wanted to analyze the effects of a parametric change of the target speed probability upon human anticipatory smooth eye movements. We have done this by re-analyzing previously collected data (see Souto et al., 2008) and by replicating and generalizing those results in a larger group of participants and different conditions, including targets with different motion direction, speed and acceleration profiles. We used target motion trajectories along both horizontal and oblique directions but kept target motion direction constant within blocks in all experiments. Second, we compared different fully predictable target kinematics, namely constant or accelerating target speeds and analyzed their effect on anticipatory eye movements. We found that anticipatory responses were strongly modulated by both constant and accelerating target speeds. We report a linear scaling of anticipatory smooth pursuit with target speed (or acceleration) probability, similar to what we previously reported for a probability bias in direction and distinct from the short-term trial history effects. We also investigated, through statistical and model-based analyses, whether the internal model of visual motion that drives anticipatory smooth pursuit would integrate information about the target’s accelerating profile. Our results, although very variable across participants, provide evidence that humans can integrate some information about acceleration and use it to anticipate the forthcoming target motion. Whether this integration is grounded on an internal, noisy representation of motion acceleration, or on an approximation of the target mean velocity over an extended temporal window remains to be further investigated.

### A linear dependence between anticipatory pursuit and the probability of target kinematic cues

In two separate experiments, we showed that the velocity of anticipatory pursuit is modulated by the constant-speed probability of visual moving targets, regardless of its fully predictable direction. These results are consistent with previous reports (as reviewed in Kowler et al., 2019) that showed that anticipatory smooth velocity is modulated by the predictability of target speed, as well as by short-term trial-sequence effects. Our results are however novel in several aspects. First, we demonstrate a parametric, linear relationship between the amplitude of the anticipatory phase and a broad range of target speed probability. Such a linear relationship is observed also over a large interval of target speeds (from 5 to above 30°/s) and is similar for targets moving along either the horizontal (**Figure 2**) or the oblique axes (**Figure 3**). In addition, similar to a previous study of our group (Damasse et al., 2018) we have provided evidence that short-term effects driven by the previous trial’s speed can coexist and interact with the long-term effects that were the main objective in the present study. In particular, experiencing a high

(low) speed trial yields an increase (decrease) of aSPv in the following trial with respect to the block’s average (**Figure 4**), but this effect is strongly modulated by the context, namely by the probability of high (or low) speed trials. Moreover, we show that higher order kinematic cues such as acceleration can also modulate eye velocity during anticipatory pursuit, again regardless of the (predictable) target motion direction and such an effect is also modulated by the probability of these higher-order kinematic cues (**Figure 5**).

Overall, these novel results extend our previous findings of a linear dependence of the anticipatory eye velocity upon the target direction probability (Damasse et al., 2018; Santos & Kowler, 2017) and further demonstrate that the statistical regularities of different motion properties are efficiently stored in memory and used to drive anticipatory visuomotor control across a timescale of several seconds to several minutes and more. Our results argue for a probabilistic coding of target velocity (direction and speed), and possibly of target acceleration as well. Yet, future work is needed to elucidate whether these aspects of target trajectories are encoded together or separately and by which neuronal populations and computational processes. It is important to notice that the present and most of the previous results demonstrating some degree of sensorimotor adaptation to the statistical regularities in the environment do not imply that probabilities are explicitly learnt and/or represented as abstract concepts in the brain. The very nature of probabilistic coding in the brain is at the heart of important lines of research, and the relation between probabilistic coding and probability-based behavior is not trivial at all. In any case, anticipatory smooth pursuit eye movements appear as an effective behavioral measure to elucidate how direction and kinematic parameters are encoded, together with their uncertainty, to control eye movements.

### Is there an internal representation of accelerating target kinematics?

In this study, we also addressed the question of *how* information about accelerating motion could be integrated in the internal model of motion that drives anticipatory eye movements. To do so, we analyzed how predictable accelerating targets affect anticipatory eye velocity compared to constant-speed targets. Target motion conditions were designed such that constant speeds can directly be compared with accelerating conditions: each value of initial speed (11, 22 or 33 °/s) was paired with different acceleration values. The linear mixed-effects regression model indicated that anticipatory eye velocity was not simply scaled based on the instantaneous initial target speed, but also on its acceleration. Moreover, the effect of acceleration increased with the initial speed (see **Figure 7**, and statistical analyses reported in the text). We performed an additional model-based analysis to tease apart the possibility that the internal model driving anticipatory smooth pursuit relies, rather than on an exact representation of target acceleration, on a simple approximation of it, namely the estimation of mean target speed across a finite temporal window close to motion onset. Leveraging on the linear relation between predictable target speed and anticipatory eye velocity, this analysis allowed us to simulate the size of the temporal window of integration and its variability across the group of participants tested in this study (**Figure 8**). Overall, the estimated temporal window of integration was strongly variable across participants and in the case of decelerating targets these estimates were often unrealistic (e.g. negative temporal windows). Interestingly, the distribution of TWI estimates with accelerating targets was clearly different from the one obtained with decelerating targets, and centered on positive and plausible values (i.e. around two-hundred milliseconds after target motion onset).

Whether and how the acceleration of moving targets is represented and used by the primate visual tracking system is still unclear (Lisberger & Movshon, 1999), even though target acceleration is a key component of many models of visual target-driven smooth pursuit eye movements (Brostek et al., 2017; Goldreich et al., 1992; Krauzlis & Lisberger, 1994). In humans, several previous studies have attempted to demonstrate a role for acceleration and whether such high-order motion cues can be learned through the history of target motion, on both the short (a few hundreds milliseconds) and medium to long (seconds to minutes and more) timescales. Bennett et al. (2007) showed that when smoothly pursuing an accelerating target which undergoes an occlusion after a short exposition (200 ms) in a random-presentation condition, human participants are not able to adaptively use the acceleration information. Instead, participants seem to store the estimate of a constant velocity and use saccades to compensate for the displacement error between the eye position and the location where the target reappears. Those authors found, however, that after a longer exposure (500-800 ms, comparable to our visual motion duration), smooth pursuit and saccades discriminate between the different acceleration profiles. Still, prediction of the target position at the end of the occlusion was not accurate. Using again the transient target-occlusion paradigm, Bennett & Barnes (2006) probed predictive smooth pursuit of accelerating targets in blocked vs random presentation conditions. They reported two interesting and complementary results: in a blocked-design paradigm (thus with highly predictable motion over a long timescale), anticipatory eye velocity occurring (1) before the target motion onset and (2) before the end of target blanking was scaled to the target acceleration. However, increasing uncertainty about target acceleration, by mixing trials, had canceled such dependency and anticipatory eye movements were no longer distinguishable between acceleration conditions. Other studies came to the same conclusion: on a short timescale acceleration information can be used to somehow control online tracking eye (and hand) movements but not to build robust predictive motor responses to moving targets, or related perceptual judgements (Kreyenmeier et al. 2022). On the other hand, predictability over a longer timescale seems to favor the integration of acceleration information for visuomotor control (Bennett & Barnes, 2006). Our results tend to corroborate the latter claim, with the *caveat* of a large inter-individual variability observed in the anticipatory behavior with predictable accelerating motion.

Early psychophysical studies have shown that the mean speed estimated over the stimulus motion duration influences the perceptual discrimination of acceleration (Brouwer et al., 2002; Gottsdanker et al., 1961; Schmerler, 1976). Watamaniuk & Heinen (2003) showed that this is also the case when judging and tracking an accelerated moving target. In addition, the duration of the temporal window during which the target kinematic information is acquired seems to influence the accuracy of acceleration estimation (Bennett et al., 2007). In the present study, our model-based analysis of the temporal window of integration highlighted a large inter-individual variability, as well as a dependence on the acceleration sign, in the timescale that would be relevant to estimate the motion of accelerating targets. Overall, several contextual factors seem to influence the encoding and processing of visual motion acceleration. The precise nature -and the mere existence!-of an explicit representation of visual acceleration in the brain remains to be elucidated.

We also need a more complete understanding of how speed and acceleration cues can be integrated through learning sensorimotor contingencies in specific tasks. A very peculiar example is the vertical tracking of a target that changes speed by following the gravity acceleration (Zago et al., 2010). What is being learned in an experimental session (e.g. probabilities of occurrence) versus the entire lifespan (e.g. the law of gravity, or friction, see Souto & Kerzel, 2013), and how this drives anticipatory pursuit responses questions the complex interactions between predictive and sensory information for an optimal tracking behavior.

Finally, in addition, in alternative or in parallel to the internal model of the retinal target speed and acceleration, reinforcement learning processes could play an important role in adapting anticipatory eye movements to predictable motion properties. Any combination of retinal position, velocity or acceleration errors could be estimated and eventually minimized over trials, akin to a *cost function*, to improve target visibility and tracking performance. Thus, we can speculate that participants could simply learn by trial and error and adapt their anticipation behavior to rapidly minimize the difference between the eyes and target position and velocity, as well as its change over time. Again, this sort of cost-minimization process remains to be thoroughly tested by future model-based experiments.

### Neuronal bases of predictive tracking and processing of different kinematic properties

Electrophysiological studies in the non-human primates have provided evidence that a small subpart of the Frontal Eye Fields (FEFsem, slightly ventral compared to the saccadic FEF) is implicated in the control of predictive smooth pursuit (e.g. Fukushima et al., 2002; MacAvoy et al., 1991, see Kowler, 2019 for a review). Darlington et al. (2018) showed that FEFsem firing rate is modulated, before visual motion onset, by the expectations about the target speed. In addition, the speed-context modulation of neuronal activity continues throughout the visually-guided phase of smooth pursuit, and it is stronger when the visual stimuli are less reliable (i.e. at lower contrast), in agreement with Bayesian integration of prior beliefs and sensory evidence. Such integration was also apparent in the oculomotor recordings, with the monkeys’ smooth pursuit eye velocity more strongly modulated by the speed context for low-contrast targets. Unfortunately, the authors could not compare the FEF preparatory activity with anticipatory eye velocity, nor did they analyze the smooth pursuit latency dependence on motion expectancy, thereby limiting the possibility to draw some correspondence with our results. A second prefrontal oculomotor field, the Supplementary Eye Fields (SEF) is also involved in the control of predictive smooth pursuit (Heinen & Liu, 1997). For instance, de Hemptinne et al. (2008) showed that the activity of a population of SEF neurons encoded the target direction expectations, as neurons became more active after the presentation of a cue indicating deterministically a target motion in the neuron’s preferred direction. The evidence for the neural substrates of predictive pursuit is much sparser in humans: Gagnon et al. (2006) have applied transcranial magnetic stimulation (TMS) pulses to the human FEFsem and SEF regions during visual tracking of sinusoidal target motion. They have reported an enhancement of predictive pursuit when TMS was applied to FEFsem at different epochs, but only in some specific conditions when TMS was applied to SEF. Several questions remain yet unanswered. First, the respective role of FEFsem and SEF in predictive eye movement is still debated. Second, how the different variables of target motion trajectories are encoded and learned is yet to be investigated. Thanks to its fast, block-designed protocol mixing different target motion cues, the present study may inspire future neurophysiological studies in non-human and human primates, focusing on the joint analysis of anticipatory responses and preparatory neural activities in these two prefrontal areas.

Our results call for a reevaluation of the role of higher-order motion cues (acceleration/deceleration) in the control and learning of predictive pursuit behavior. There is very little evidence that the primate nervous system encodes visual acceleration explicitly, in the visual or in the oculomotor systems. Lisberger & Movshon (1999) measured MT single neurons’ responses to image acceleration, but did not find evidence that those neurons’ activity varied with acceleration. They found, however, that the simulated nonlinear readout of a population of MT neurons was correlated to image acceleration (though not to its deceleration). Similarly, Price et al. (2005) found speed tuning in MT single neurons, but not an acceleration or deceleration tuning. However, Schlack et al. (2007) showed that a linear classifier can extract acceleration signals from the MT population response, given that the MT neurons’ tuning to speed depended on the acceleration and deceleration contexts of the task. Note, however, that these earlier studies focused mainly on primate area MT while other parietal (MST) and frontal (FEF) cortical areas might contribute to represent complex target motion trajectories and higher-order kinematics. Future work shall elucidate how position, velocity and acceleration cues are jointly or independently encoded across the visuo-oculomotor distributed network, to represent and learn target trajectories for the efficient control of action.

## Conclusion

In this study, we showed that when the target speed is predictable, human participants show a linear dependence of anticipatory eye velocity upon the speed probability that is comparable to the dependence found for target direction probability. Moreover, participants also show anticipatory responses adjusted to accelerating target kinematics, and to their probability across-trials. Overall, this study contributes to the broad existing literature about the sensory and cognitive control of eye movements by better characterizing the role of predictive information about the target kinematics.

## Data availability statement

The data and analysis scripts are available (DOI 10.17605/OSF.IO/SYD3T).

## Acknowledgements

This work was funded by the *Agence Nationale de la Recherche* (ANR-PREDICTEYE 18-CE37--0019 to GSM, AM) and the *Fondation pour la Recherche Médicale* (Equipe FRM 2018 to GSM) and by Aix-Marseille Université (PhD Doctoral extension funding to VCM). We would like to thank Mr Alexis Ulian for helping in the data collection of the Experiment 2A as part of his Master internship.

## Supplementary material

### Final models for the LMM analysis

Exp 1A:

*aSPv ∼ 1 + P(HS) + (1 + P(HS) | participant)*

*aSPv ∼ 1 + P(HS)*Tv_N-1_ + (1 + P(HS) +Tv_N-1_ | participant)*

Exp 1B, constant speed probability-mixtures:

*aSPv* ∼ *1 + P(v33) + axis + (1 + P(v33) + axis | participant)*

*aSPv ∼ 1 + P(v33)*Tv_N-1_ + (1 + P(v33) +Tv_N-1_ | participant)*

Exp 2A-B, accelerating target probability-mixtures:

*aSPv ∼ 1 + prob + axis + exp + axis:exp + (1 + prob + axis | participant)*

Exp 3, comparison between fully predictable blocks:

*aSPv ∼ 1 + v0*accel + (1 + v0 | participant)*

Exp3, categorical model for pairwise comparisons:

*aSpv ∼ 1 + condition + (1 | participant)*

### LMM analysis - result tables

Exp1A: Anticipatory Parameters
|  | *Dependent variable:* |
| --- | --- |
|  | aSPv |
| P(HS) | 3.488<sup>***</sup> (3.138, 3.838)<br>t = 19.541<br>p = 0.000 |
| Constant | 2.759<sup>***</sup> (2.036, 3.482)<br>t = 7.480<br>p = 0.000 |
| Random Effects |  |
| Groups | 3 |
| sd(Constant) | 0.77 |
| sd(P(HS)) | 0.33 |
| *Note:* * p < 0.01; ** p < 0.001; *** p < 1e-04 |  |

**Exp 1A: Anticipatory Parameters– Sequential Effects**
| *Dependent variable:* |  |
| --- | --- |
|  | aSPv |
| Tv<sub>N-1</sub>[LS] | -0.501<sup>***</sup> (-0.663, -0.340)<br>t = -6.073<br>p = 0.000 |
| P(HS) | 4.357<sup>***</sup> (3.983, 4.732)<br>t = 22.827<br>p = 0.000 |
| P(HS):Tv<sub>N-1</sub>[LS] | -2.744<sup>***</sup> (-3.102, -2.386)<br>t = -15.030<br>p = 0.000 |
| Constant | 4.427<sup>***</sup> (3.801, 5.053)<br>t = 13.853<br>p = 0.000 |
| Random Effects |  |
| Groups | 3 |
| sd(Constant) | 0.70 |
| sd(P(HS)) | 0.10 |
| sd(Tv<sub>N-1</sub>[LS]) | 0.29 |

*Note:* \* p < 0.01; \*\* p < 0.001; \*\*\* p < 1e-04

Exp 1B: Anticipatory Parameters – Const Speed
| *Dependent variable:* |  |
| --- | --- |
|  | aSPv |
| P(v33) | 2.743<sup>***</sup> (1.824, 3.663)<br>t = 5.847<br>p = 0.000 |
| Axis[vert.] | -0.740 (-1.464, -0.016)<br>t = -2.002<br>p = 0.046 |
| P(v33):Axis[vert.] | -0.674<sup>***</sup> (-0.985, -0.363)<br>t = -4.252<br>p = 0.00003 |
| Constant | 2.707<sup>***</sup> (1.740, 3.673)<br>t = 5.490<br>p = 0.00000 |
| Random Effects |  |
| Groups | 13 |
| sd(Constant) | 1.82 |
| sd(P(v33)) | 1.68 |
| sd(Axis[vert.]) | 1.32 |

*Note:* \* p < 0.01; \*\* p < 0.001; \*\*\* p < 1e-04

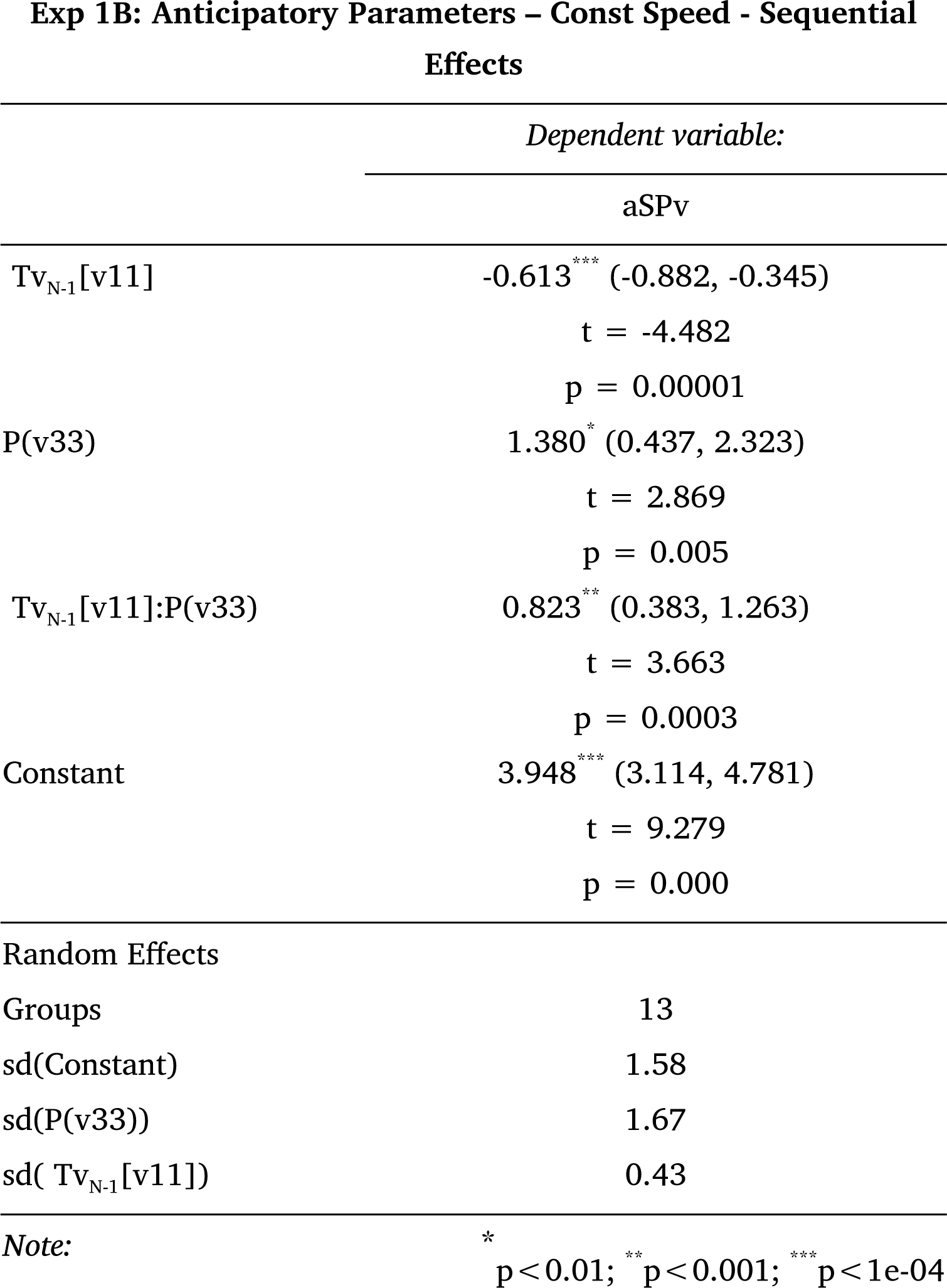

Exp 2A-B: Anticipatory Parameters – Accelerating Target
|  | Dependent variable: |
| --- | --- |
|  | aSPv |
| P(vdec) | 1.884<sup>***</sup> (1.111, 2.657)<br>t = 4.779<br>p = 0.00001 |
| Axis[vert.] | -0.460 (-1.264, 0.343)<br>t = -1.123<br>p = 0.262 |
| Exp.[const.Time] | 1.683<sup>***</sup> (1.410, 1.956)<br>t = 12.086<br>p = 0.000 |
| Exp.[const.Time]:Axis[vert.] | -1.651<sup>***</sup> (-2.027, -1.275)<br>t = -8.611<br>p = 0.000 |
| Constant | 3.041<sup>***</sup> (2.099, 3.983)<br>t = 6.326<br>p = 0.000 |
| Random Effects |  |
| Groups | 16 |
| sd(Constant) | 1.96 |
| sd(P(V3)) | 1.58 |
| sd(Axis[vert.]) | 1.66 |
| Note: | * p < 0.01; ** p < 0.001; *** p < 1e-04 |

**Exp 3: Anticipatory Parameters – Parametric Variables**
| *Dependent variable:* |  |
| --- | --- |
|  | aSPv |
| V0 | 0.246<sup>***</sup> (0.149, 0.343)<br>t = 4.963<br>p = 0.00000 |
| Accel | 0.017<sup>*</sup> (0.004, 0.030)<br>t = 2.640<br>p = 0.009 |
| V0:Accel | 0.001<sup>*</sup> (0.0004, 0.002)<br>t = 2.905<br>p = 0.004 |
| Constant | 0.364 (-0.324, 1.052)<br>t = 1.037<br>p = 0.300 |
| Random Effects |  |
| Groups | 7 |
| sd(Constant) | 0.95 |
| sd(v0) | 0.14 |
| Groups | 7 |
| Note: | <sup>*</sup> p < 0.01; <sup>**</sup> p < 0.001; <sup>***</sup> p < 1e-04 |

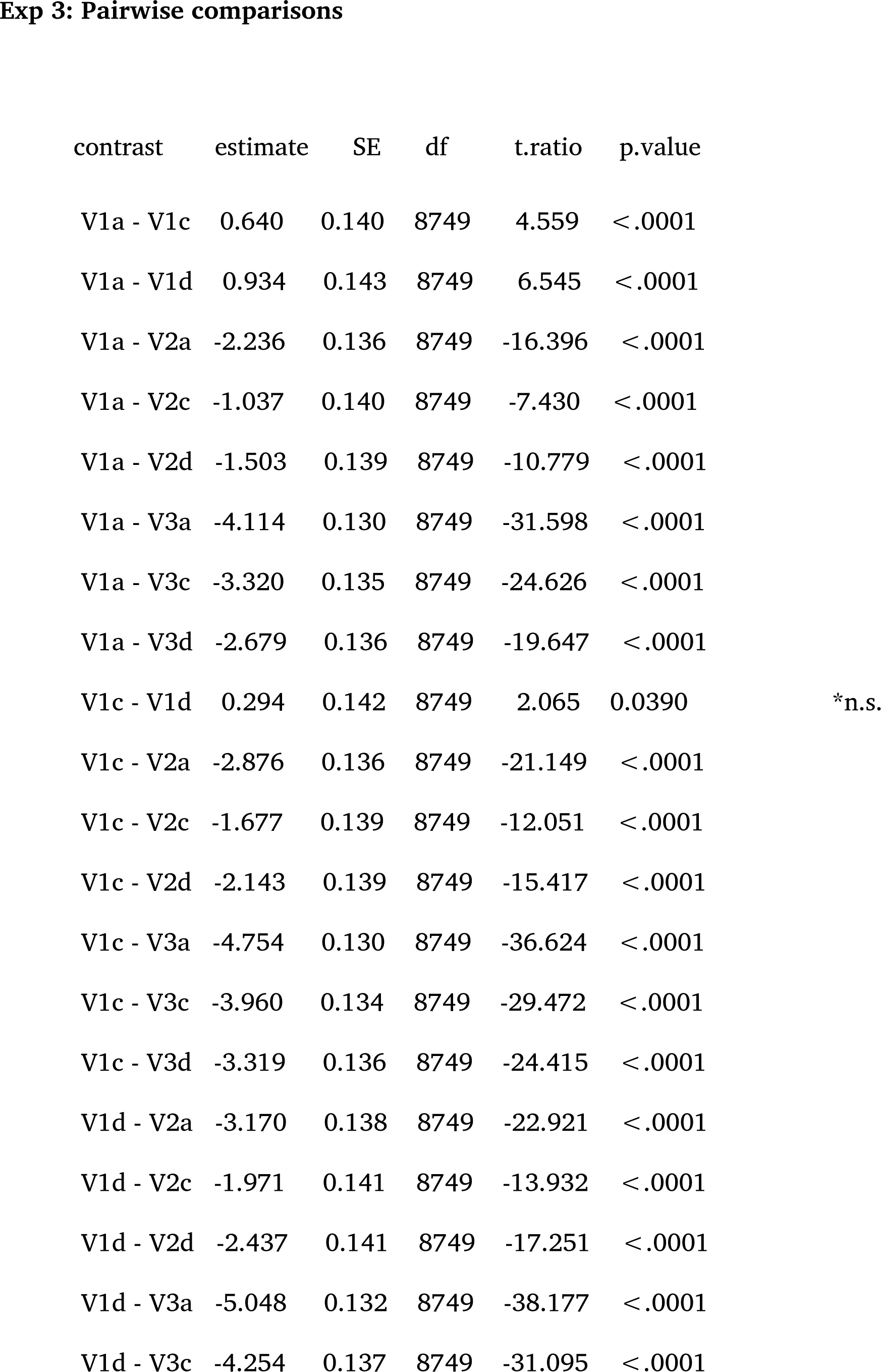
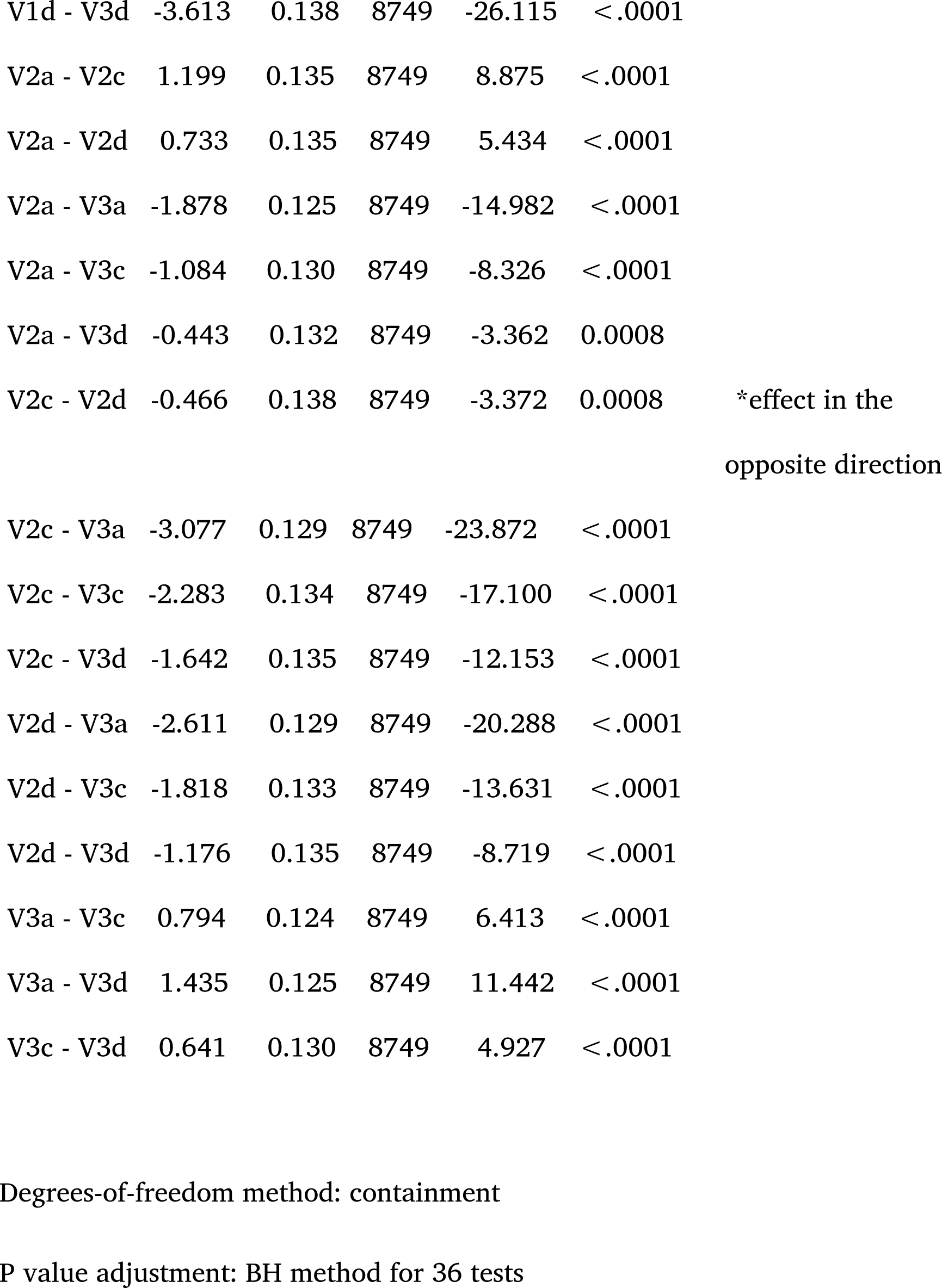

## Notes

### Competing Interest Statement

The authors have declared no competing interest.

### Summary of Updates

The manuscript now focuses on anticipatory smooth pursuit (aSP) velocity and includes one extra experiment to test directly the effects of acceleration on aSP

